# Automated Harmonization and Large-Scale Integration of Heterogeneous Biomedical Sample Metadata Using Large Language Models

**DOI:** 10.1101/2024.10.26.620145

**Authors:** Koichi Higashi, Zenichi Nakagawa, Takuji Yamada, Hiroshi Mori

**Affiliations:** Genome Evolution Laboratory, National Institute of Genetics, Mishima, Shizuoka 411-8540, Japan; School of Life Science and Technology, Institute of Science Tokyo, Ookayama, Tokyo 152-8550 Japan; Metagen, Inc., 246-2 Mizukami, Kakuganji, Tsuruoka, Yamagata, 997-0052, Japan; Metagen Theurapeutics, Inc., 246-2 Mizukami, Kakuganji, Tsuruoka, Yamagata, 997-0052, Japan; digzyme, Inc., Toranomon, Minato-ku, Tokyo, 105-0001, Japan; Genome Diversity Laboratory, National Institute of Genetics, Mishima, Shizuoka 411-8540, Japan

**Keywords:** Microbiome, Database, Large Language Models, Metadata

## Abstract

The exponential growth of biomedical data has created an urgent need for efficient integration and analysis of heterogeneous sample metadata across studies. However, current methods for harmonizing and standardizing these metadata are largely manual, time-consuming, and prone to inconsistencies. Here, we present a novel computational framework that leverages large language models (LLMs) to automate the harmonization and large-scale integration of diverse biomedical sample metadata. Our approach combines semantic clustering techniques with LLM-driven natural language processing to extract, interpret, and standardize metadata from various sources, including research papers, supplementary tables, and text data from public databases. We demonstrate the efficacy of our framework by applying it to thousands of human gut microbiome papers, successfully extracting and integrating metadata from over 400,000 samples. Our method achieved a 50% recovery rate of manually curated metadata, significantly outperforming traditional rule-based methods. Furthermore, our framework enabled the creation of a unified, searchable database of standardized metadata, facilitating cross-study analyses and revealing previously obscured patterns in microbiome composition across diverse populations and conditions. The scalability and adaptability of our approach suggest its potential applicability to a wide range of biomedical fields, potentially accelerating meta-analyses and fostering new insights from existing data. This work represents a significant advancement in biomedical data integration, offering a powerful tool for researchers to unlock the full potential of accumulated scientific knowledge.

## Introduction

The field of biomedical research has experienced an unprecedented increase in data generation over the past two decades, driven by advancements in high-throughput technologies and decreasing costs of data acquisition[1]. This data deluge has been particularly pronounced in microbiome research[2], where next-generation sequencing technologies have enabled researchers to investigate the complex microbial communities inhabiting various environments, including the human body. The human gut microbiome, in particular, has emerged as a crucial factor in understanding human health and disease[3–6].

As the volume of biomedical data continues to expand exponentially, the scientific community faces a significant challenge: how to effectively integrate, analyze, and derive meaningful insights from this wealth of information[7,8]. Central to this challenge is the critical role of metadata – the contextual information that describes the conditions under which biological samples were collected, processed, and analyzed[9–11]. In microbiome research, metadata encompasses a wide range of factors, including host demographics, environmental conditions, dietary habits, medical history, and experimental protocols[12,13]. This information is crucial for interpreting sequencing data, identifying patterns, and drawing valid conclusions across studies.

The importance of metadata in microbiome research cannot be overstated[12]. It provides the necessary context for understanding the complex interactions between microorganisms and their environment. For instance, host factors such as age[14], diet[15], antibiotic use[16], and geographical location[17,18] can significantly influence the composition of microbial communities in the human gut. Without accurate and comprehensive metadata, researchers risk drawing incomplete or misleading conclusions from their analyses. Moreover, the integration of metadata across multiple studies is essential for meta-analyses[13,19], which can reveal broader patterns and insights not apparent in individual studies.

However, the current state of metadata in biomedical research, particularly in microbiome studies, is far from ideal. Despite efforts to standardize metadata reporting, such as the Minimum Information about any (x) Sequence (MIxS) guidelines proposed by the Genomic Standards Consortium[20], there remains a lack of consistency in how metadata is recorded, formatted, and shared across different studies[12,21,22]. This heterogeneity poses significant obstacles to data integration and cross-study comparisons. Researchers often encounter incompatible formats, inconsistent terminologies, and varying levels of detail in metadata from different sources. The process of harmonizing and standardizing this diverse metadata is typically manual, time-consuming, and prone to errors, creating a bottleneck in the research pipeline[23,24].

The challenges of metadata harmonization are further compounded by the sheer volume of published research. With thousands of microbiome studies published annually[25], manually curating and standardizing metadata across all these studies becomes an increasingly daunting task. This situation not only hampers the progress of individual research projects but also impedes the field’s ability to leverage the full potential of accumulated data for generating new hypotheses and insights.

Recent advancements in artificial intelligence, particularly in the domain of natural language processing (NLP) and large language models (LLMs)[26–29], offer promising avenues for addressing these challenges. LLMs, trained on a vast corpora of text, have demonstrated remarkable capabilities in understanding context, extracting information, and generating human-like text[30,31]. These models have the potential to revolutionize how we approach metadata extraction, harmonization, and integration in biomedical research.

In this study, we present a novel computational framework that leverages the power of LLMs to automate the process of harmonizing and integrating heterogeneous biomedical sample metadata. Our approach combines advanced NLP techniques with semantic clustering to extract, interpret, and standardize metadata from diverse sources, including research papers, supplementary tables, and text data from public databases. By applying this framework to a large corpus of human gut microbiome studies, we demonstrate its efficacy in creating a unified, standardized metadata resource that facilitates cross-study analyses and uncovers previously obscured patterns in microbiome composition across diverse populations and conditions.

## Results

We developed EMBERS (Encompassing Microbiome-Bibliome Extraction and Retrieval System), a novel computational framework for automated harmonization and large-scale integration of heterogeneous biomedical sample metadata. EMBERS was applied to a corpus of 26,435 human gut microbiome-related research papers, demonstrating its efficacy in extracting, harmonizing, and analyzing metadata at an unprecedented scale.

### EMBERS Framework Overview and Performance

The EMBERS framework consists of two main components: EMBERS-MINE for extracting metadata from individual papers, and EMBERS-FUSE for integrating and harmonizing metadata across studies (Figure 1 and Figure 2). Throughout both stages, we strategically combined traditional computational methods with targeted use of LLMs to address specific challenges, while maintaining overall control through a custom Python program.

**Figure 1.**
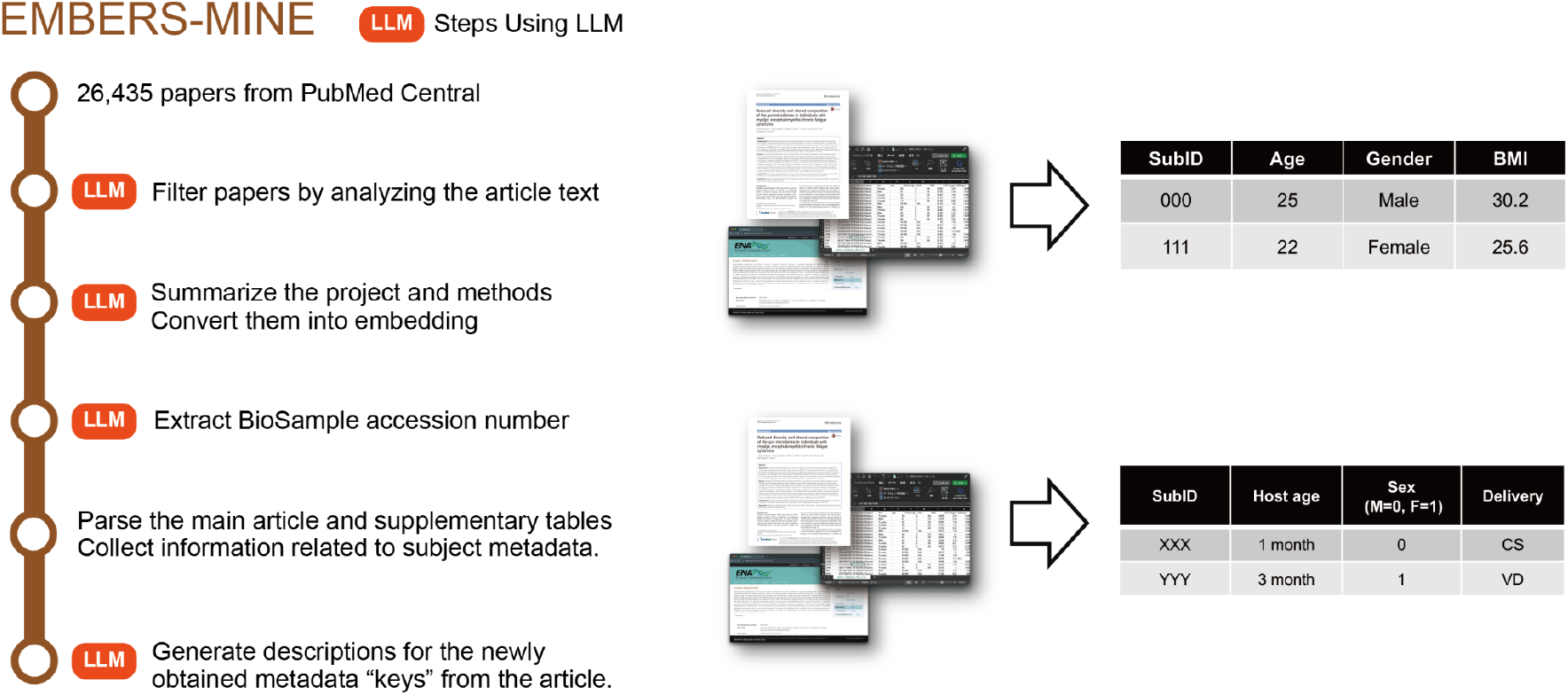
Overview of Sample Metadata Extraction from Individual Papers

**Figure 2.**
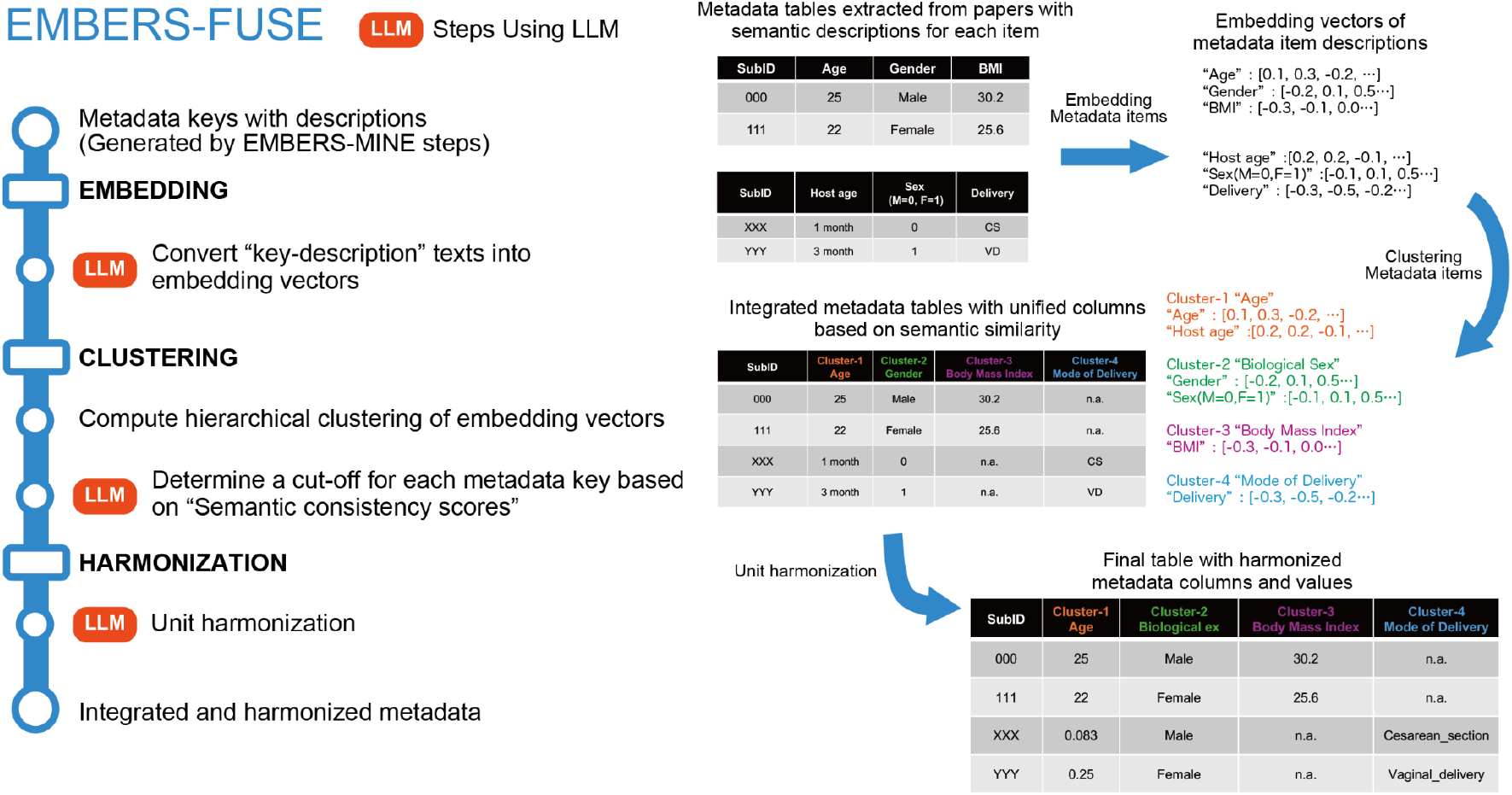
Overview of Integration and Harmonization of Data Across All Papers

Each paper processed through EMBERS-MINE (Figure 1) undergoes three key steps:

1. Initial paper assessment using LLMs to verify relevance to original human gut microbiome research and distinguish from meta-analyses or non-relevant studies
2. Extraction of structured metadata from supplementary materials and main text using specialized parsers for different file formats
3. Context-aware interpretation of extracted metadata through LLM-driven analysis, generating semantic descriptions that capture the full meaning of each metadata item within the paper’s context

The resulting paper-specific metadata then flows into EMBERS-FUSE (Figure 2), which performs:

1. Vector embedding generation for metadata descriptions using specialized language models optimized for semantic similarity
2. Novel semantic clustering to group related metadata across different studies, allowing for identification of equivalent concepts despite varying terminology
3. Automated unit harmonization through LLM-generated conversion scripts, ensuring consistency across studies
4. Integration into a unified, queryable database

EMBERS-MINE successfully processed the entire corpus of 26,435 papers, extracting relevant metadata from both main text and supplementary materials. The extraction process covered key metadata categories including age, biological sex, BMI (Body Mass Index), and mode of delivery, among others. EMBERS-FUSE then integrated this extracted metadata, resulting in a harmonized dataset comprising information from over 400,000 samples.

### Key Concepts

#### 1. LLM-Enhanced Metadata Extraction (EMBERS-MINE)

Traditional rule-based methods struggle with the diversity and context-dependency of metadata representations in scientific literature. LLMs to overcome these challenges. A key innovation in EMBERS-MINE is the use of LLMs for context-aware metadata interpretation. The system generates descriptive text for each metadata item based on the paper’s full context, providing semantic understanding beyond simple keyword matching. This step is crucial for the subsequent integration in EMBERS-FUSE, as it enables more accurate categorization of diverse metadata representations across studies.

EMBERS-MINE employs LLMs to assess paper relevance, effectively distinguishing original research from meta-analyses or non-relevant studies. This context-aware filtering significantly improved the quality of input data for metadata extraction. As a supplementary benefit, our LLM-based approach also improved the accuracy of distinguishing between the study’s original samples and referenced external datasets when extracting BioSample accession numbers, a task that traditional regex-based methods often struggle with due to lack of contextual understanding.

#### 2. Semantic Clustering for Metadata Integration (EMBERS-FUSE)

The challenge in metadata integration lies in the inconsistent naming and representation of similar concepts across different studies. For instance, “Gender” in one study might be labeled as “Sex(M=0, F=1)” in another. To address this, we developed a novel “semantic clustering” approach in EMBERS-FUSE, combining hierarchical clustering of embedding vectors with LLM-based semantic consistency evaluation.

This method effectively groups diverse metadata representations into coherent semantic categories, enabling meaningful integration across studies. By leveraging the semantic understanding provided by LLMs, our approach can identify and group conceptually similar metadata items despite surface-level differences in naming or formatting. The details of this clustering algorithm are further elaborated in the Methods section.

#### 3. Adaptive Unit Harmonization

Metadata tables often use inconsistent unit systems, such as representing age in years in one study and months in another. Direct LLM-based conversion for each value is computationally infeasible for large datasets and exceeds token limits. Our solution to this challenge is an innovative approach that balances the need for context-aware conversion with computational efficiency.

For each study, we use LLMs to generate Python code for unit conversion based on the study’s specific context. This code is then executed within EMBERS to perform efficient, study-specific unit standardization. This approach allows for scalable processing of large datasets while maintaining the flexibility to handle diverse and complex unit representations across different studies.

### Accuracy Evaluation

We evaluated EMBERS using a manually curated “ground truth” dataset from 100 papers, containing 22,104 samples and 49,712 metadata items. The evaluation focused on metadata recovery efficiency (Recall) and accuracy of recovered metadata (Precision), comparing against a baseline of MIxS term-based extraction (See Methods).

EMBERS achieved an average recall of approximately 50% (Figure 3A), significantly outperforming MIxS-based extraction. Interestingly, recovery rates were often bimodal, with many papers achieving either near 100% or 0% recall (black dots in Figure 3A show Per-Paper Performance). EMBERS consistently improved recall for papers where MIxS-based extraction failed completely, demonstrating its robustness across diverse document structures and writing styles.

**Figure 3.**
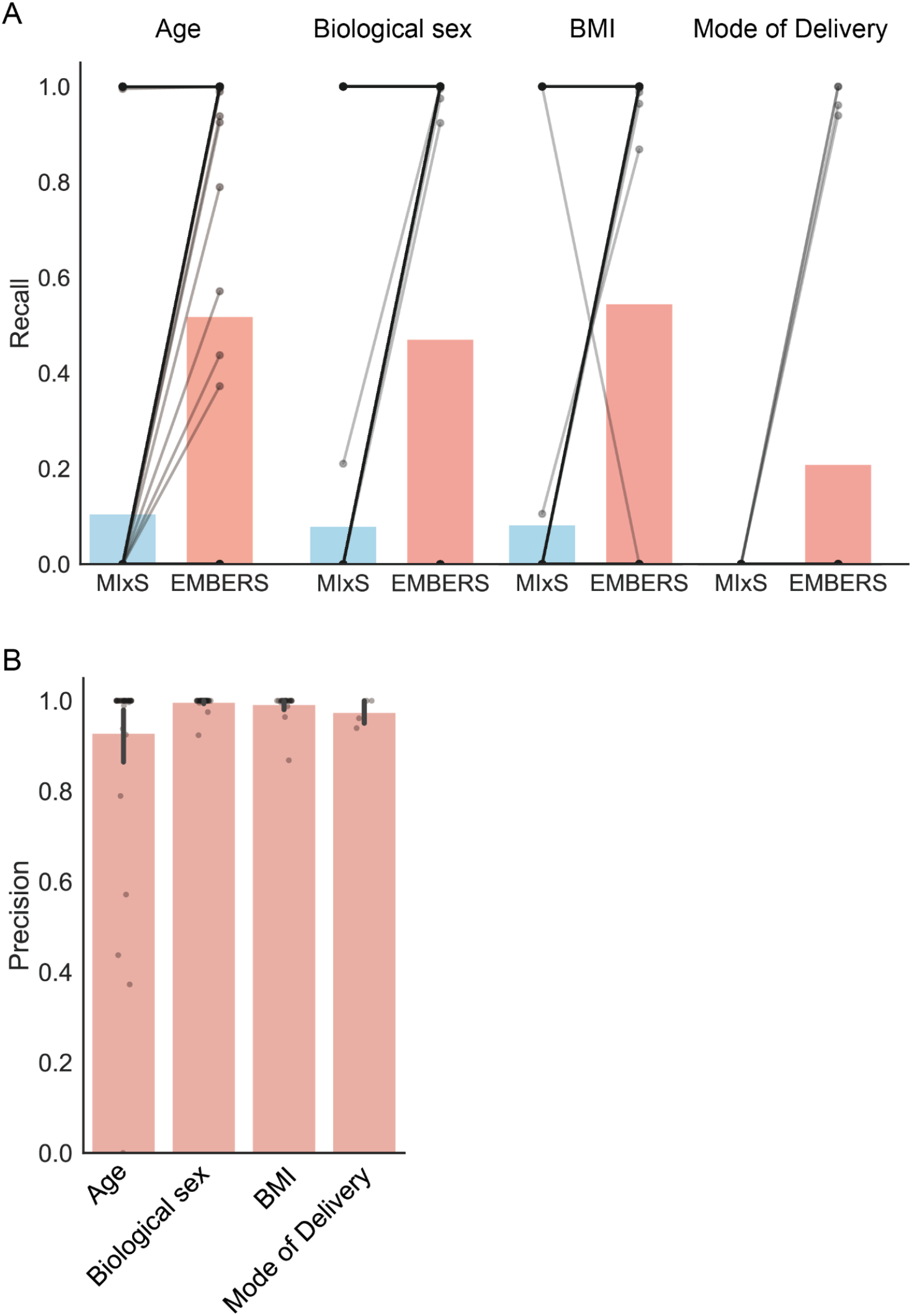
Precision and recall evaluation of metadata extraction for Age, Biological sex, BMI (Body Mass Index), and Mode of Delivery. (A) Comparison of recall between MIxS term-based extraction and our EMBERS approach. Black dots connected by lines represent individual paper results, while bar graphs indicate overall recall for each metadata type. (B) Precision of metadata extracted using EMBERS. Black dots denote precision for individual papers, and bar graphs show overall precision for each metadata category.

Across all metadata categories, precision was consistently near 100% (Figure 3B), indicating high reliability of extracted metadata. This suggests that while EMBERS may occasionally fail to extract certain metadata items, the items it does extract are highly accurate.

One notable challenge was observed with the “Mode of Delivery” metadata, which showed exceptionally low recall. This is likely due to the complexity of linking this information between infant and maternal metadata across different tables, highlighting an area for future improvement in our framework.

### Analysis of Harmonized Metadata

The large-scale integration of metadata enabled novel insights into the landscape of human gut microbiome research. Our analysis revealed a trimodal distribution in subject ages across studies (Figure 4A). We observed a high concentration of studies focusing on subjects under 1 year old, followed by a second peak in the 20-30 age range, and a third peak around 60 years of age. This distribution likely reflects distinct research focuses: infant development studies, research related to women’s pregnancy and childbirth, and investigations into age-related diseases in the elderly population, respectively.

**Figure 4.**
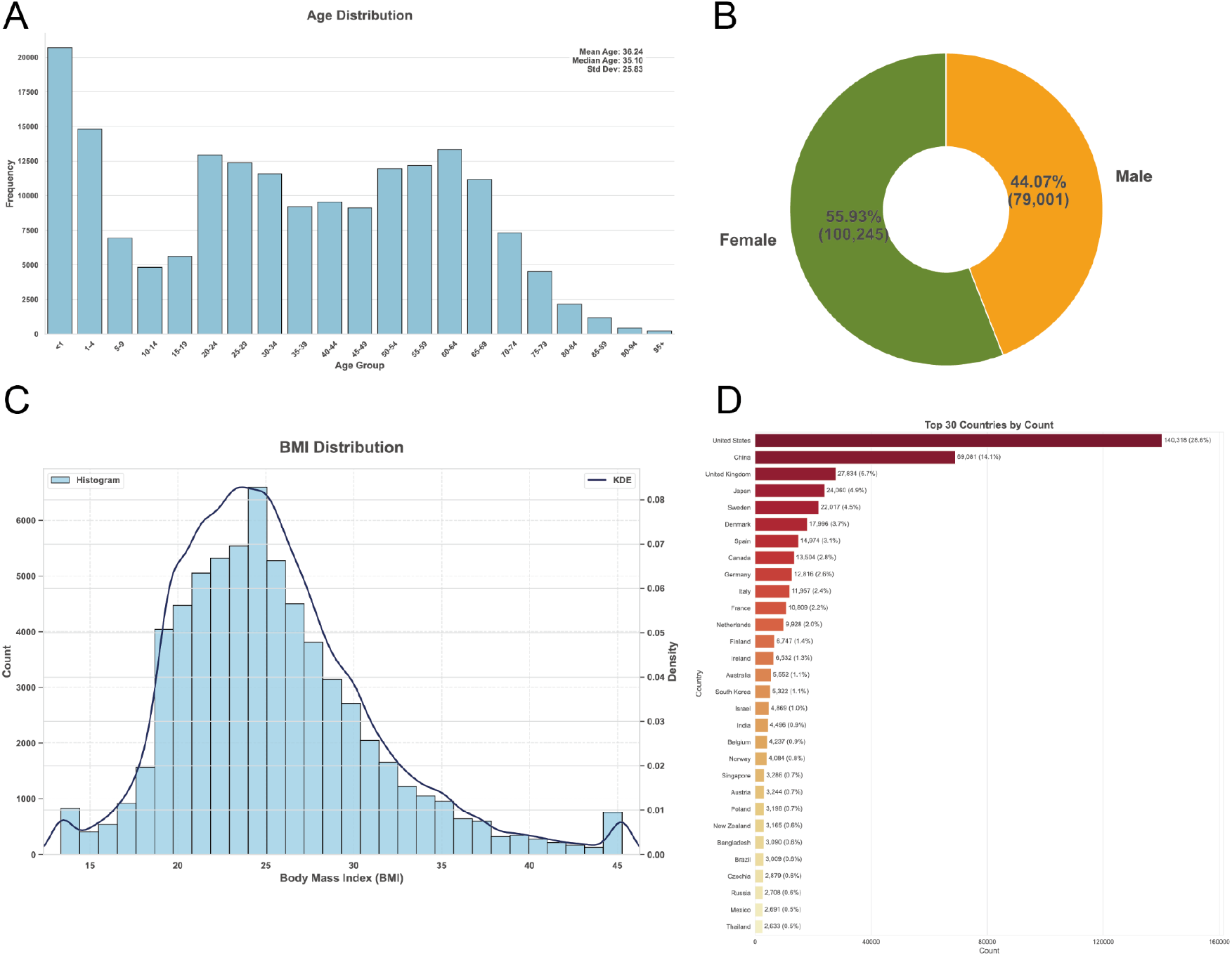
Distribution of extracted metadata from human gut microbiome studies. (A) Age distribution. (B) Biological sex distribution. (C) Body Mass Index (BMI) distribution. (D) Geographical distribution by country. Each panel represents the distribution of successfully extracted metadata for the respective category using the EMBERS approach.

Examination of the BMI distribution (Figure 4C) showed a peak around 25, which is slightly shifted to the right compared to global patterns[32], but not dramatically so. Interestingly, we observed slightly higher proportions of subjects at the extreme ends of the BMI spectrum - both those with very low BMI (< 15) and those with very high BMI (> 40). This suggests that while the majority of studies focus on populations with BMIs in the normal to overweight range, there is also targeted research on individuals with extreme body compositions.

Further analysis of the harmonized data uncovered a disproportionate representation of certain geographical regions, highlighting potential gaps in global microbiome research coverage (Figure 4D)[33]. Since some of the population-based gut microbiome studies do not submit the microbiome metadata to the BioSample database and only submit to controlled access databases (e.g., Database of Genotypes and Phenotypes (dbGaP), European Genome-phenome Archive (EGA), and Japanese Genotype-phenotype Archive (JGA))[34–36], these geographic region statistics may not accurately reflect study trends. Additionally, examination of biological sex data revealed a slight overrepresentation of female subjects in the studied cohorts (Figure 4B). The overrepresentation of female subjects may reflect the trend of infant and maternal gut microbiome studies[37].

### Application of Harmonized Metadata

To demonstrate the utility of our harmonized metadata database, we conducted a large-scale analysis of relationships between metadata and microbiome composition patterns. The taxonomic composition data was obtained from the Sandpiper dataset[38], which provides large-scale taxonomic composition data from shotgun metagenome samples analyzed using the SingleM method[39]. We constructed a database linking taxonomic compositions to metadata for 40,465 shotgun metagenome samples that overlapped with our extracted metadata tables.

A UMAP (Uniform Manifold Approximation and Projection)[40] visualization of taxonomic compositions, color-coded by various metadata categories, revealed complex associations between host factors and microbial community structures (Figure 5A). This visualization, representing an unprecedented scale of integrated microbiome and metadata analysis, underscores the power of our harmonized database in uncovering previously obscured relationships in microbiome data.

**Figure 5.**
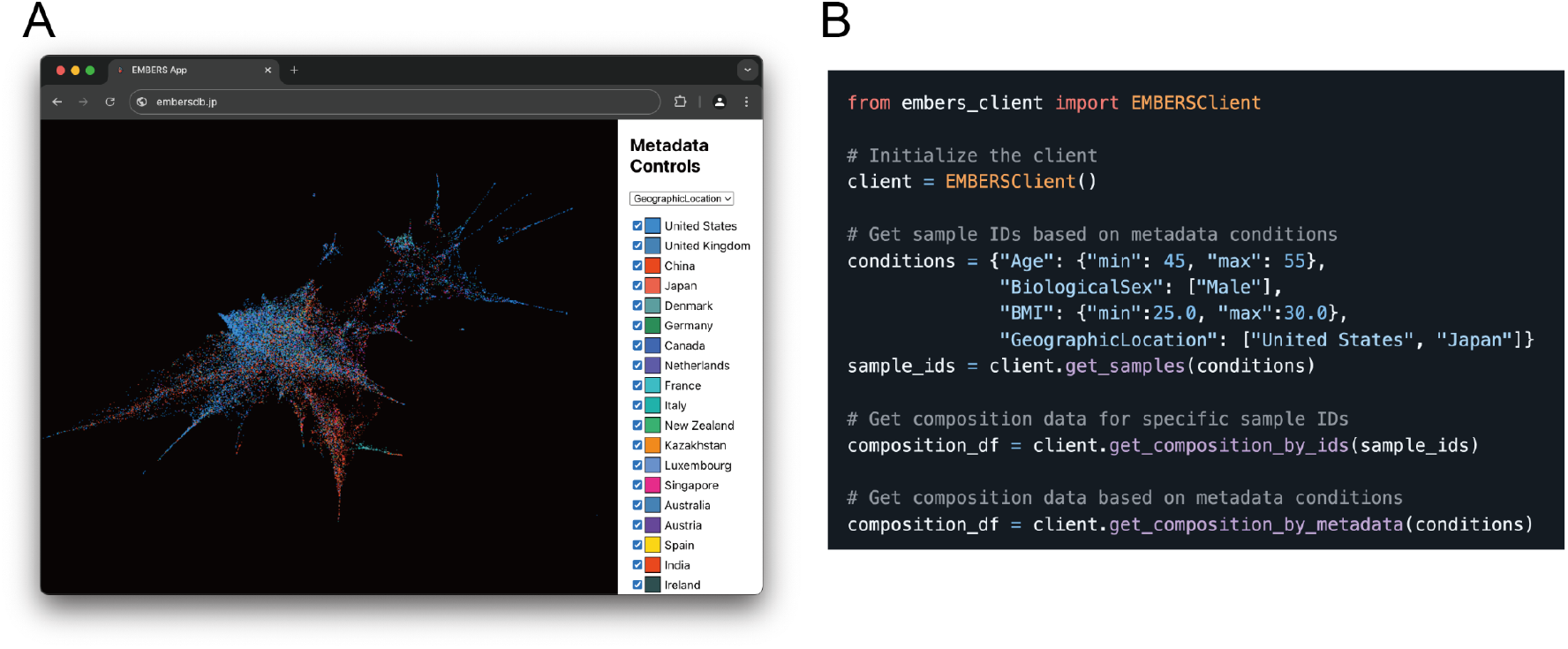
Visualization and access tools for integrated human gut microbiome metadata and taxonomic composition data. (A) Screenshot of an interactive web application displaying the distribution of metadata extracted by EMBERS for human gut microbiome studies. This visualization tool allows users to explore taxonomic composition data from shotgun metagenomic analyses in relation to various extracted metadata categories. (B) Example usage of the EMBERS-CLIENT Python package, which facilitates retrieval of sample IDs and taxonomic composition data based on subject attributes from the integrated metadata database. This client application enables researchers to efficiently query and extract relevant data subsets for downstream analyses.

To facilitate broader use of this harmonized dataset, we developed EMBERS-CLIENT, a Python package that allows researchers to easily query the database and retrieve relevant sample sets (Figure 5B). This tool enables users to obtain Pandas DataFrames of gut microbiome taxonomic compositions based on specific subject metadata criteria, significantly simplifying large-scale, metadata-driven analyses in microbiome research.

In summary, EMBERS has demonstrated its capability to efficiently extract, harmonize, and integrate metadata from a large corpus of biomedical literature. The resulting harmonized database, coupled with user-friendly tools for data access and visualization, provides a valuable resource for the microbiome research community, enabling large-scale, metadata-driven analyses and potentially accelerating discoveries in this rapidly evolving field.

## Discussion

In this study, we have developed and implemented a novel computational framework for the automated harmonization and large-scale integration of heterogeneous biomedical sample metadata, with a specific focus on human gut microbiome research. Our approach, which strategically combines LLMs with traditional computational methods, demonstrates significant advancements in the field of metadata extraction and integration. The creation of a comprehensive, harmonized database of microbiome metadata, along with the development of the EMBERS-CLIENT Python package, represents a substantial contribution to the research community, enabling unprecedented opportunities for cross-study analyses and meta-studies.

The superior performance of our method compared to MIxS-based extraction underscores the potential of integrating advanced AI capabilities with established computational techniques. Achieving a 50% recovery rate of manually curated metadata represents a significant improvement over the metadata retrieval method based on MIxS terms, which are often not fully adhered to by researchers[12]. However, this figure also highlights the complexity of the task and the substantial room for enhancement that remains.

A critical aspect of our framework is the strategic use of LLMs. While these models played a significant role in addressing specific challenges throughout our pipeline, it is crucial to emphasize that the entire process was orchestrated and controlled by a custom Python program. This program invoked OpenAI’s ChatCompletion and TextEmbedding APIs only when necessary, ensuring efficient use of computational resources and maintaining overall control of the workflow. This approach allowed us to harness the power of LLMs without relying on them exclusively, striking a balance between advanced AI capabilities and traditional computational methods.

The strategic limitation of LLM usage to specific, critical tasks mitigated the risk of hallucinations – a common concern with LLMs[41,42]. By primarily utilizing LLMs for tasks that did not heavily rely on domain-specific knowledge, we minimized the potential for significant errors. However, it is important to acknowledge that the integration of LLMs with existing methods can introduce new types of errors. For instance, in our metadata clustering results, we occasionally observed semantically divergent items grouped together (e.g., “Sexual orientation” included in the “Biological sex” cluster). These instances, while rare, highlight the need for ongoing refinement and human oversight in AI-assisted data processing.

Our analysis revealed several challenging scenarios that contributed to the gap between our automated extraction and manual curation:

1. Embedded Metadata in Sample IDs: The embedding of crucial metadata within sample identifiers requires a more nuanced understanding of study-specific naming conventions.
2. Cross-Paper Information: Our current framework, designed to process papers individually, struggles with metadata that can only be fully recovered by cross-referencing multiple papers.
3. Non-Standard Reporting Formats: Despite standardization efforts, many papers present metadata in highly customized or study-specific formats, posing significant challenges for automated extraction.
4. Implicit Information: Capturing metadata that is implied rather than explicitly stated remains a challenge for our automated system, underscoring the continued value of human expertise in interpreting complex scientific information.

These challenges highlight areas for future improvement and research directions. Enhancing our system’s ability to interpret complex sample naming conventions, developing methods for cross-document analysis, improving the handling of non-standard formats, and incorporating more sophisticated domain knowledge inference could significantly boost the recovery rate and expand the applicability of our framework.

The creation of a large-scale, harmonized metadata database and the EMBERS-CLIENT Python package opens up new avenues for microbiome research. The ability to easily query and analyze metadata across numerous studies can facilitate the discovery of patterns and relationships previously obscured by data heterogeneity. However, it is crucial for researchers to be aware of the potential biases in the compiled data, as our analysis of metadata distributions reveals patterns specific to subjects in gut microbiome studies rather than general population demographics.

The rapidly evolving landscape of human gut microbiome research, with thousands of new studies published annually, underscores the need for continuous updates and expansions of our EMBERS framework. Large-scale cohort studies, which are becoming increasingly common[4,43], further emphasize the limitations of manual data integration approaches. Recognizing this, we have designed EMBERS with scalability and ease of update in mind. The modular nature of our system, particularly the separation of EMBERS-MINE for individual paper processing and EMBERS-FUSE for overall integration, facilitates the incorporation of new research data. While EMBERS-FUSE requires a complete rerun for comprehensive integration, the paper-specific nature of EMBERS-MINE allows for efficient processing of newly added studies.

We are committed to maintaining the relevance and comprehensiveness of our database through regular updates. These updates will not only incorporate new research but also refine our extraction and integration methodologies based on emerging patterns in data reporting. Furthermore, we envision expanding the application of EMBERS beyond the human gut microbiome. Given the similarities in subject metadata reporting patterns, we anticipate a relatively straightforward adaptation of our framework to other human microbiome niches, such as the oral cavity and skin. This expansion will provide a more holistic view of human-associated microbial communities and their interactions.

Looking further ahead, we are exploring the potential application of EMBERS to environmental microbiome studies, including soil and marine ecosystems. However, we acknowledge that this expansion presents unique challenges. Environmental metadata tends to be more diverse and complex compared to human subject data, often embedded within the text of papers rather than in structured tables. Addressing these challenges will likely require more sophisticated use of LLMs for information extraction from unstructured text. This adaptation process will not only enhance the capabilities of EMBERS but also contribute to the broader field of automated scientific literature analysis.

In conclusion, our work represents a significant step forward in addressing the metadata challenge in biomedical research, particularly in the field of microbiome studies. By demonstrating the effective integration of LLMs with traditional computational methods, we have paved the way for more sophisticated approaches to data harmonization and integration. As we continue to refine our methods and address the identified challenges, we anticipate further improvements in metadata extraction and integration capabilities. This ongoing work has the potential to greatly enhance our ability to leverage existing data, facilitate more comprehensive meta-analyses, and ultimately accelerate scientific discovery in the microbiome field and beyond.

## METHODS

### Data Collection

Our study utilized a comprehensive dataset of 26,435 human gut microbiome research papers. These papers were identified through a systematic search of PubMed using the query “human gut microbiome” OR “human gut metagenome”. From the initial search results, we selected papers that were available in full text through PubMed Central, ensuring accessibility for our automated analysis pipeline.

### EMBERS Framework Overview

The EMBERS (Encompassing Microbiome-Bibliome Extraction and Retrieval System) framework was developed to address the challenge of extracting and harmonizing metadata from a large corpus of biomedical literature. EMBERS consists of two primary components: EMBERS-MINE and EMBERS-FUSE.

EMBERS-MINE focuses on the extraction and interpretation of metadata from individual papers, leveraging advanced natural language processing techniques and large language models. EMBERS-FUSE, on the other hand, is responsible for the integration and harmonization of metadata across all processed papers, employing novel semantic clustering approaches and unit standardization methods.

The framework operates in a sequential manner, with EMBERS-MINE processing each paper individually, followed by EMBERS-FUSE integrating the extracted metadata across the entire corpus. This modular design allows for scalability and flexibility in processing large volumes of biomedical literature. The entire codebase for these processes is publicly available at https://github.com/khigashi1987/EMBERS.

### EMBERS-MINE: Sample Metadata Extraction

The EMBERS-MINE component of our framework was designed to extract and interpret metadata from individual research papers.

### Language Model and API Usage

Throughout the EMBERS-MINE process, we utilized OpenAI’s API services. Specifically, we employed the gpt-4-turbo model[29] via the ChatCompletion API for text generation and interpretation tasks. For text embedding, we primarily used the text-embedding-3-large model through the Text Embedding API, with one exception noted below.

### Paper Filtering and Relevance Assessment

To ensure the relevance of papers in our dataset, we implemented an LLM-based text analysis approach. The gpt-4-turbo model was prompted to assess each paper’s abstract and determine its relevance to original human fecal metagenome studies. Papers identified as meta-analyses or non-relevant were excluded from further processing.

### Research Landscape Visualization

To provide a comprehensive view of the human gut microbiome research landscape, we implemented two additional analyses:

1. Study Summary Embedding: We used the ChatCompletion API to generate one-sentence summaries of each paper. These summaries were then embedded using the Text Embedding API and visualized using UMAP[40] to reveal sub-genres within the field.
2. Experimental Process Embedding: Recognizing the potential for experimental bias in microbiome studies, we segmented method descriptions into individual experimental steps. Each step was embedded into a high-dimensional vector, allowing for quantitative comparison of experimental approaches across studies.

The results have been implemented as an interactive web application, allowing users to freely explore the findings. The application is accessible at http://embersdb.jp.

### BioSample Accession Number Extraction

We developed a hybrid approach combining regular expressions with LLM-based context analysis for extracting BioSample accession numbers. Regular expressions were used to identify potential accession number patterns, while the gpt-4-turbo model was prompted to analyze the surrounding context and determine whether the identified numbers were indeed related to the study’s original samples.

### Metadata Table Parsing

A custom parser was developed to handle the diverse formats of supplementary tables in Excel format. This parser was specifically designed for parsing metadata tables in supplementary files provided in Excel. For tables embedded in the main text and supplementary data in PDF format, the Camelot Python package (v0.11.0, https://camelot-py.readthedocs.io/) was employed to accurately extract and parse the table data.

### Metadata Semantic Interpretation

To enhance the interpretability of extracted metadata, we utilized the gpt-4-turbo model to generate descriptive text for each metadata item. The model was prompted to analyze the full text of each paper and provide semantic descriptions that captured the context and meaning of each extracted metadata item.

### EMBERS-FUSE: Data Integration and Harmonization

The EMBERS-FUSE component was developed to integrate and harmonize metadata across all processed papers.

### Metadata Embedding

For generating vector representations of semantic descriptions, we primarily used the text-embedding-3-large model. However, for embedding metadata item names specifically, we found that the text-embedding-ada-002 model produced superior results and thus was used for this particular task.

### Semantic Clustering

We developed a novel “semantic clustering” approach to address the challenges of creating meaningful metadata item clusters. This method combines hierarchical clustering of embedding vectors with LLM-based semantic consistency evaluation. The process is as follows:

1. Initial hierarchical clustering of embedding vectors using cosine similarity as the distance metric and average linkage method.
2. Breadth-first traversal of the resulting cluster tree, starting from the root.
3. At each node, we calculate a Semantic Consistency Score (SCS) using the following procedure:
  i. Due to the large number of metadata items in each node (especially near the top of the hierarchy), we employ a strategic sampling method: First, we apply Principal Component Analysis (PCA) to reduce the dimensionality of the embedding space for the metadata items in the current node. We then perform K-means clustering on this reduced space. From each resulting K-means cluster, we sample metadata items evenly, ensuring a broad representation of the embedding space.
  ii. We randomly select 100 metadata items using this stratified sampling approach.
  iii. These 100 items are then presented to the gpt-4-turbo model, which is prompted to assess whether all items belong to the same semantic category.
  iv. The gpt-4-turbo model provides a Semantic Consistency Score between 0.0 and 1.0, reflecting the degree of semantic similarity among the sampled items.
4. If the SCS exceeds a predefined threshold of 0.9, we define the current node as a semantic cluster and terminate exploration of its sub-nodes.
5. If the SCS is below the threshold, we continue the breadth-first traversal to the node’s children.

This approach balances the need for semantic coherence with computational efficiency, minimizing the number of API calls to the language model while ensuring a comprehensive exploration of the metadata item space. The stratified sampling method helps mitigate potential bias that could arise from densely clustered regions in the embedding space, ensuring that the SCS calculation considers the full range of semantic variability within each node.

### Unit Harmonization

To address the complex challenge of unit conversion across diverse metadata representations, we developed an LLM-driven approach:

1. For each study, we collect a representative set of metadata item names and value examples.
2. The gpt-4-turbo model is prompted to generate Python code capable of converting these values to a standardized unit system.
3. The generated Python code is evaluated to perform the actual conversions.

This method allows for flexible handling of various unit representations, including cases where conversion rules are embedded within metadata item names (e.g., “Sex(M=0,F=1)”).

### Manual curation process

The extraction of metadata embedded in the papers was conducted manually. Specifically, 3,000 papers were initially selected through keyword searches in PubMed, from which 100 papers containing metagenomic and sample metadata were manually selected. These selected papers primarily included metadata in supplemental data, where sample IDs and corresponding metadata were provided in PDF or Excel formats. The extracted data were organized into text format. Additionally, metadata item names and units were standardized by consulting the main text of each paper. Ultimately, from the 100 papers, 22,104 samples and 49,712 metadata items with values (15,593 for “Age”, 19,817 for “Biological Sex”, 7,373 for “Body Mass Index” and 6,929 for “Mode of Delivery”) were extracted and prepared as a ground truth set for validating metadata extraction by our method.

### Accuracy Evaluation

To assess the performance of our automated metadata extraction and harmonization framework, we conducted a comprehensive evaluation using a manually curated “ground truth” dataset described above.

We evaluated our framework’s performance using standard metrics of precision and recall:

Precision: The proportion of correctly extracted metadata items among all extracted items.

Recall: The proportion of correctly extracted metadata items among all relevant items in the ground truth dataset.

For each metadata category (Age, Biological Sex, BMI, and Mode of Delivery), we calculated precision and recall on a per-paper basis and then computed the overall metrics across all papers.

As a baseline for comparison, we also evaluated the performance of a method using the Minimum Information about any (x) Sequence (MIxS) terms[20], as proposed by the Genomic Standards Consortium (GSC). This comparison allowed us to quantify the improvement our approach offers over existing standardized terminologies. The MIxS terms used are included in the human-gut extension package and are as follows: host_age (MIXS:0000255), host_sex (MIXS:0000811), and host_body_mass_index (MIXS:0000317). For mode of delivery, which lacks a corresponding term in the MIxS standard, we provisionally used “host_delivery” as a search term to enable comparison, although this represents a departure from strict MIxS-based extraction. To account for minor variations in MIxS terms representations, we implemented a normalization process for both the extracted metadata and the ground truth data. This process involved standardizing spaces, underscores, and letter cases to ensure fair comparison.

### Application Development and Data Integration

To complement our metadata integration efforts with corresponding microbial taxonomic composition data, we utilized the Sandpiper database v0.3.0[38]. Sandpiper provides taxonomic composition information for large-scale shotgun metagenomic public data, analyzed using the SingleM method[39]. We downloaded the Sandpiper database and extracted sample IDs that overlapped with our integrated metadata dataset. For these overlapping samples, we retrieved the genus-level taxonomic composition information and reconstructed it into a taxonomic composition table. This integration process resulted in a final dataset comprising 40,465 samples with both metadata and taxonomic composition information.

To facilitate efficient access and querying of this large-scale dataset, we developed a web API that manages sample metadata using SQLite3 RDBMS (version 3.45.3) and stores taxonomic composition data in a Chroma vector database (version 0.5.13). This API allows for searching and extracting samples based on metadata criteria, enabling researchers to perform complex queries across the integrated dataset.

We provide two primary means of accessing the integrated data. First, we developed an interactive web application (Figure 5A) that implements a 3D embedding visualization of the taxonomic composition data for 40,465 samples using UMAP. This application utilizes Three.js (r169) for high-performance rendering of the 3D visualization, allowing users to explore the data interactively with options to filter and color-code samples based on various metadata attributes. Second, we created a Python client package (Figure 5B) that allows programmatic access to the database. This client enables users to query the database and retrieve relevant sample sets based on specific metadata criteria, providing functionality to obtain Pandas DataFrames of gut microbiome taxonomic compositions for further analysis in Python environments.

## Data availability

The tools developed in this study are publicly available. EMBERS, a suite of programs used for extracting subject metadata from the PubMed Central article dataset, is accessible at https://github.com/khigashi1987/EMBERS. To facilitate local access to the integrated subject metadata database, retrieve sample IDs, and obtain corresponding microbial taxonomic composition tables, we have developed EMBERS-CLIENT, a Python package available at https://github.com/khigashi1987/EMBERS_CLIENT. Additionally, a web application for visualizing taxonomic compositions can be accessed at http://embersdb.jp.

## Acknowledgments

We are grateful to Dr. Ken Kurokawa (National Institute of Genetics, Japan) for his insightful discussions and valuable suggestions that contributed to the development of this research. This work was supported by JSPS KAKENHI Grant Number JP22H04925 (PAGS) and Japan Science and Technology Agency (DICP JPMJND2206).

## Author contributions

K.H., T.Y., and H.M. conceptualized the study. K.H. developed the methodology and software, and performed data visualization. Z.N. conducted the extensive manual data curation. K.H. and Z.N. collaborated on the validation process. K.H., T.Y., and H.M. wrote the original draft of the manuscript and were responsible for its review and editing. T.Y. and H.M. administered the project, acquired funding, and provided overall supervision. All authors have read and agreed to the published version of the manuscript.

## Competing interests

The authors declare no competing interests.

T.Y. is a founder of Metagen Inc., Metagen Therapeutics Inc., and digzyme Inc. Metagen Inc. focuses on the design and control of the gut environment for human health. Metagen Therapeutics Inc. focuses on drug discovery and development which utilizes microbiome science. All of the companies had no control over the interpretation, writing, or publication of this work.

## Notes

http://embersdb.jp/

